# Fluoxetine improves behavioural deficits induced by chronic alcohol addiction by alleviating RNA editing of 5-HT_2C_ receptors in astrocytes

**DOI:** 10.1101/751065

**Authors:** Zexiong Li, Shanshan Liang, Shuai Li, Beina Chen, Manman Zhang, Maosheng Xia, Dawei Guan, Alexei Verkhratsky, Baoman Li

**Affiliations:** Practical Teaching Centre, School of Forensic Medicine, China Medical University, Shenyang, People’s Republic of China.; Department of Orthopaedics, The First Hospital, China Medical University, Shenyang, People’s Republic of China.; Department of Forensic Pathology, School of Forensic Medicine, China Medical University, Shenyang, China.; Faculty of Biology, Medicine and Health, The University of Manchester, Manchester, UK.; Department of Poison Analysis, School of Forensic Medicine, China Medical University, Shenyang, China.

**Keywords:** 5-HT2C receptor, chronic alcohol addiction, ADAR2, ATP, astrocytes

## Abstract

The alcoholism and major depressive disorder are common comorbidity, with alcohol-induced depressive symptoms being eased by selective serotonin re-uptake inhibitors (SSRIs), although the mechanisms underlying pathology and therapy are poorly understood. Chronic alcohol consumption affects the activity of serotonin 2C receptors (5-HT_2C_R) by regulating adenosine deaminases acting on RNA (ADARs) in neurones. Astrogliopathic changes contribute to alcohol addiction, while decreased release of ATP from astrocytes can trigger depressive-like behaviours in mice. In this study, we discovered that chronic alcohol addiction increased editing of RNA of 5-HT_2C_R via up-regulating the expression of ADAR2, consequnetly reducing the release of ATP from astrocytes induced by 5-HT_2C_R agonist, MK212. At the same time SSRI antidepressant fluoxetine decreased the expression of ADAR2 through the transactivation of EGFR/PI3K/AKT/cFos signalling pathway. Reduction in ADAR2 activity eliminated the RNA editing of 5-HT_2C_R *in vivo* and increased release of astroglial ATP which was suppressed by chronic alcohol consumption. Meanwhile, fluoxetine improved the behavioural and motor symptoms induced by alcohol addiction and decreased the alcohol intake. Our study suggests that the astrocytic 5-HT_2C_R contribute to alcohol addiction; fluoxetine thus can be used to alleviate depression, treat alcohol addiction and improve motor coordination.

## Introduction

Alcohol abuse, characterized by excessive drinking and inability to control alcohol consumption, reflecting serious psychological dependence, is a widespread addictive disorder^1^. Concomitant occurrence of alcoholism and major depressive disorder (MDD) is common, while symptoms of depression induced by alcohol abuse can gradually disappear with the alcohol abstinence^2^. The mechanism(s) underlying the role of alcohol addiction in formation of depressive symptoms is poorly understood. Clinical management of alcohol-induced depressive symptoms mostly relies on selective serotonin re-uptake inhibitors (SSRIs)^3^, with the therapeutic effects developing in 3-4 weeks after therapy commencement; the relevant pharmacological mechanisms remain unclear.

Serotonin 2C receptors (5-HT_2C_R), are Gq-protein coupled receptors widely expressed in neurones and astrocytes in the central nervous system (CNS). After chronic exposure to alcohol, the activity of 5-HT_2C_R can be modified by adenosine deaminases acting on RNA (ADARs), which convert adenosine (A) to inosine (I) at five sites of 5-HT_2C_R mRNA; this process is associated with the expression of 24 potential edited isoforms^4, 5^. Aberrant activity of 5-HT_2C_R was suggested to link to the alcohol addiction and the neuropsychiatric disorders^6, 7^. The mRNA editing of neuronal 5-HT_2C_R in mice can be also triggered by stress as well as by exposure to SSRIs^8^. Alcohol abuse, however, can affect not only neuronal networks but also act on astrocytes, which are primarily responsible for homeostasis and catabolism of central neurotransmitters^9–11^. According to our previous studies, dysfunction of astrocytes contributes to pathophysiology of MDD, while SSRI antidepressant fluoxetine acts as an agonist of 5-HT2BR in astrocytes; this action being linked to fluoxetine anti-depressive activity^12–16^. Fluoxetine selectively increases the expression of 5-HT2BR in astrocytes, and up-regulates 5-HT_2C_R expression in neurones in conditions of chronic unpredictable mild stress (CUMS)^17^. In astrocytes, fluoxetine increases RNA editing of 5-HT2BR through ADAR2 signalling cascade^17–19^.

Recent studies demonstrated that decreased secretion of ATP from astrocytes can induce the depressive-like behaviours in mice, while fluoxetine exposure was shown to increase the level of extracellular ATP in rodents^20, 21^. Astrocytes can release ATP in response to physiological stimulation^22^, and hence we analysed the relationship between astroglial ATP release and editing of serotonin receptors in the context of alcohol addiction and fluoxetine therapy.

## Materials and methods

### Animals

The CD-1 mice were purchased from Charles River, Beijing, China. For *in vitro* experiments, newborn mice were used for making primary cultures. For animal experiments, 10 - 12 weeks old mice weighting about 25 g were used; animals were raised in standard housing conditions (22 ± 1℃; light/dark cycle of 12/12h), with water and food available *ad libitum*. All experiments were performed in accordance with the US National Institutes of Health Guide for the Care and Use of Laboratory Animals (NIH Publication No. 8023) and its 1978 revision, and all experimental protocols were approved by the Institutional Animal Care and Use Committee of China Medical University, No. [2019]059.

### Materials

Primary antibodies against ADAR2, 5-HT_2C_R, β-actin and cFos, were purchased from Santa-Cruz Biotechnology (Santa, CA, USA). Secondary anti-body for western blot, anti-mouse IgG HRP conjugated, was purchased from Promega (Madison, WI, USA). Fluor-conjugated secondary anti-body was purchased from Abcam (Cambridge, MA). Most chemicals, including MK212 (2-Chloro-6-(1-piperazinyl)-pyrazine hydrochloride), SB20474 (N-(1-Methyl-1H-5-indolyl)-N’-(3-methyl-5-isothiazolyl) urea), LY294002(2-(4-Morpholinyl)-8-phenyl-1(4H)-benzopyran-4-one hydrochloride), PP1 (4-amino-5-(4-methylphenyl)-7-(t-butyl)pyrazolo-d-3,4-pyrimidine) and AG1478 (N-[(2R)-2-(hydroxamidocarbonymethyl)-4-methylpentanoyl]-Ltryptophan methylamide), were purchased from Sigma (St Louis, MO, USA). ATP assay kits were purchased from Beyotime Biotechnology (Shanghai, China). TUNEL cells apoptosis assay kits were purchased from Roche (Mannheim, Germany).

### Alcohol addiction model and drug treatment

As described previoulsy^23^, 24 mice were randomly divided into two groups to be fed either with alcohol or water. Mice in alcohol addiction model group were exposed to gradually increasing alcohol concentrations: 3% for 5 days, 7% for 5 days, 11% for 5 days, 15% for 7 days and 20% for 7 days. Alcohol solution was only water source for alcohol group. After this protocol, alcohol preference test was performed. Subsequently, mice feed with water or alcohol were further randomly divided into two groups with intraperitoneal injection (i.p.) of fluoxetine or normal saline (NS) for one week after which behavioral tests were performed.

### Primary culture of astrocytes

As described previously^14, 24^, newborn mice were used to isolate astrocytes. The neopallia of cerebral hemispheres were isolated, dissociated and filtered. Isolated astrocytes were grown in Dulbecco’s Minimum Essential Medium (DMEM) with 7.5 mM glucose supplemented with 20% horse serum. Astrocytes were incubated at 37 ℃ in a humidified atmosphere of CO_2_/air (5:95%).

### Western blotting

As described previoulsy^12^, protein concentrations of samples were detected by Lorry method, with bovine serum albumin as the standard. In brief, each sample contained 50 μg protein, which were applied on slab gels of 10% polyacrylamide. After transfer to nitrocellulose membranes, TBS-T (Tris Buffered Saline with Tween 20) containing 5% skimmed milk powder was used to block samples for 2 h. Then, the primary antibodies was utilized to incubate the nitrocellulose membranes for 2 h, which were specific to 5HT2CR, cFos and ADAR2. Samples were incubated for another 2h with the corresponding secondary-antibodies. Staining was visualized by ECL detection reagent, and images were acquired with an electrophoresis gel imaging analysis system.

### PCR sequencing

As described previoulsy^12, 23^, polymerase chain reaction amplification was performed with 5-HT_2C_R primers, forward (5’ agatatttgtgccccgtctgg 3’) and reverse (5’ aagaatgccacgaaggaccc 3’). The amplified samples were then used for cDNA sequence test. Complementary DNA sequencing was carried out by TaKaRa biotechnology Co. Ltd, and RNA editing efficiencies were calculated by the peak heights.

### ATP content assay

The culture medium was collected after incubating cells with or without 4 mg/ml alcohol and 1 μM fluoxetine for 24 h; subsequnetly cells were treated with 30 μM MK212 (a selective agonist of 5-HT_2C_R) or NS (normal saline) for 15 min,. As described previoulsy^26^, ATP assay kit (Beyotime Biotechnology, Shanghai, China) was used to detect ATP concentration. ATP standard samples were prepared with same culture medium. 20 μl samples or standard ATP samples were added into opaque 96-well plates. 100 μl ATP assay reagent were added into 96-well plates to mix with samples for 5 min. Chemoluminescence intensity was detected by fluorescence reader (Infinite M200 Pro, Tecan, Switzerland).

### Behavioural tests

#### Alcohol preference test

As described previoulsly^23^, alcohol drinking percent indicates alcohol preference. After 12 h of food and water deprivation, mice were provided with two pre-weighed bottles, including one bottle that contained 20 % alcohol and a second bottle filled with water, for 2 h. The percentage preference was calculated according to the following formula: % preference = (alcohol intake/(alcohol + water intake)) × 100 %.

#### Open field test

The open field test is an anxiety-based test, as previously described^27^. The mice were placed in the centre square of an open field box (60 × 60 × 40 cm) divided into nine squares, and behaviours were recorded for 6 min. The parameters used for analysis included the total travel distance and time spent in the central area.

#### Tail suspension test

As described previously^27^, comparing the immobility time in tail suspension test reflects despair-like behaviour. Mice were suspended by its tail at 20 cm from ground for 6 min. The time of immobility was recorded to calculate percentage.

#### Rotating rod test

As per our previous description^27^, time of staying on rotating rod indicates motion balance ability. Each mouse was placed on a rotating bar, which was set to a rotation speed of up to 20 rpm during the test. The time spent on the rotating bar was recorded as the latent period. The latency before falling was recorded using a stopwatch, with a maximum of 90 s.

### Hematoxylin-eosin (HE) staining

After fixation with 4% paraformaldehyde, cells were washed twice by double distilled water for 5 min. Subsequently cells were stained with hematoxylin solution for 5 min and 1% eosin solution for 3 min. The number of cells was imaged by a microscope (Axio Scan Z1, Zeiss, Germany).

### TUNEL staining

As described previoulsy^28^, cultured astrocytes were fixed with 4% paraformaldehyde. The cells were permeabilized with 4% bovine serum albumin for 1 hr. Apoptosis in the astrocytes was detected via TUNEL assays (cell death detection; Roche, Mannheim, Germany), according to the manufacturer’s protocol. Then, the astrocytes were stained for the astrocytic marker GFAP followed by incubation with DAPI solution at 1:1000. Images were captured with a confocal scanning microscope (DMi8, Leica, Germany). The apoptotic cells were expressed as a percentage of the TUNEL-positive cells among the DAPI-positive cells.

### Statistics

The differences among multiple groups were analyzed by one-way or two-way analysis of variance (ANOVA) followed by a Tukey post hoc multiple comparison test for unequal replications using SPSS 20.0 software. All statistical data in the text are presented as the mean ± SEM, the value of significance was set at p < 0.05.

## Results

### Alcohol increases expression of cFos and ADAR2 cultured in astrocytes

As compared with control group, the protein expression of cFos was increased by 112.50 ± 24.71% at 4 mg/ml (n = 6, p = 0.020) and by 171.07 ± 44.53% at 8 mg/ml (n = 6, p = 0.021) (Fig. 1B), the protein level of ADAR2 was also up-regulated by 131.72 ± 20.36% (n = 6, p = 0.008) and by 289.42 ± 33.64% (n = 6, p = 0.002) at these two doses of alcohol (Fig. 1C). Based on these results, 4 mg/ml alcohol was used for the subsequent experiments, as this concentration of alcohol did not evoke apoptosis. As shown in Fig. 1D and E, alcohol exposure did not change the number of astrocytes stained with HE and observed in the bright field. In the TUNEL assay, the ratio of TUNEL+/DAPI+ in alcohol group was 86.02 ± 22.76% of control group (n = 6, p = 0.715; Fig. 1G), this difference however did not reach statistical significance.

**Figure 1.**
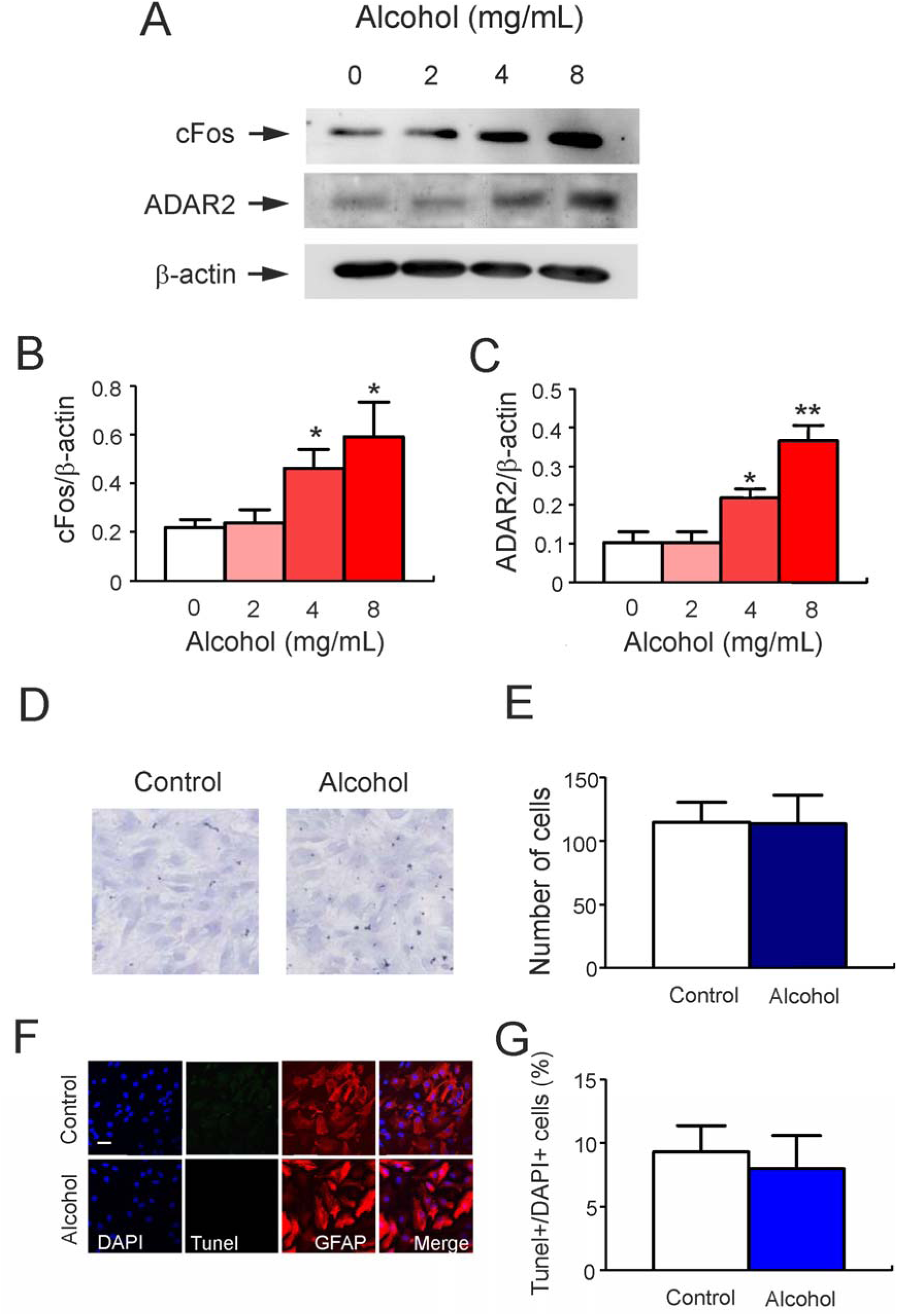
The dose response of alcohol on the expression of cFos and ADAR2 in astrocytes. **(A-C)** Primary cultured astrocytes were incubated in absence of alcohol (0 mg/ml as control group, which was treated with normal saline) or in the presence of 2, 4 or 8 mg/ml alcohol for 24 hours. **(A)** Representative western blots showing protein levels of cFos, ADAR2 and β-actin (a housekeeping protein). Similar results were obtained from six independent experiments. Average expression of cFos or ADAR2 was quantitated as ratio between cFos and β-actin (**B**), or ratio between ADAR2 and β-actin (**C**). Data are shown as mean ± SEM, n = 6. *Indicates statistically significant (p<0.05) difference from control group; **indicates statistically significant (p<0.05) difference from any other group. **(D and E)** Primary cultured astrocytes were incubated in presence of normal saline (control group) or 4 mg/mL alcohol for 24 h. **(D)** Representative images showing HE staining. **(E)** The average number of cells is presented as mean ± SEM, n = 6. **(F)** Apoptosis as detected by the TUNEL assay. Astrocytes were stained with GFAP (red), and cell nuclei were immunostained with DAPI (blue). Bar 20 μm. **(G)** Percentage of cell death was determined by the ratio of TUNEL+ and DAPI+ cells. Data are represented as mean ± SEM, n = 6.

### Fluoxetine suppresses expression of cFos and ADAR2 in cultured astrocytes

Exposure of primary cultured astrocytes to 1 μM fluoxetine decreased expression of both cFos and ADAR2 to 47.36 ± 4.38% (n = 6, p = 0.022) and 58.38 ± 3.51% (n = 6, p = 0.002) respectively (compared to the control; Figs. 2C and 2D). After treatment of cultures with 4 mg/ml alcohol for 1 day, the protein expression of cFos in alcohol plus fluoxetine group was decreased to 83.40 ± 2.11% of alcohol plus NS group (n = 6, p = 0. 030) (Fig. 2C), and the protein expression of ADAR2 in alcohol plus fluoxetine group was eliminated to 83.12 ± 1.84% of alcohol plus NS group (n = 6, p = 0.009) (Fig. 2D). Conversely, the treatment with fluoxetine with or without alcohol did not change the protein expression of 5-HT_2C_R in astrocytes.

**Figure 2.**
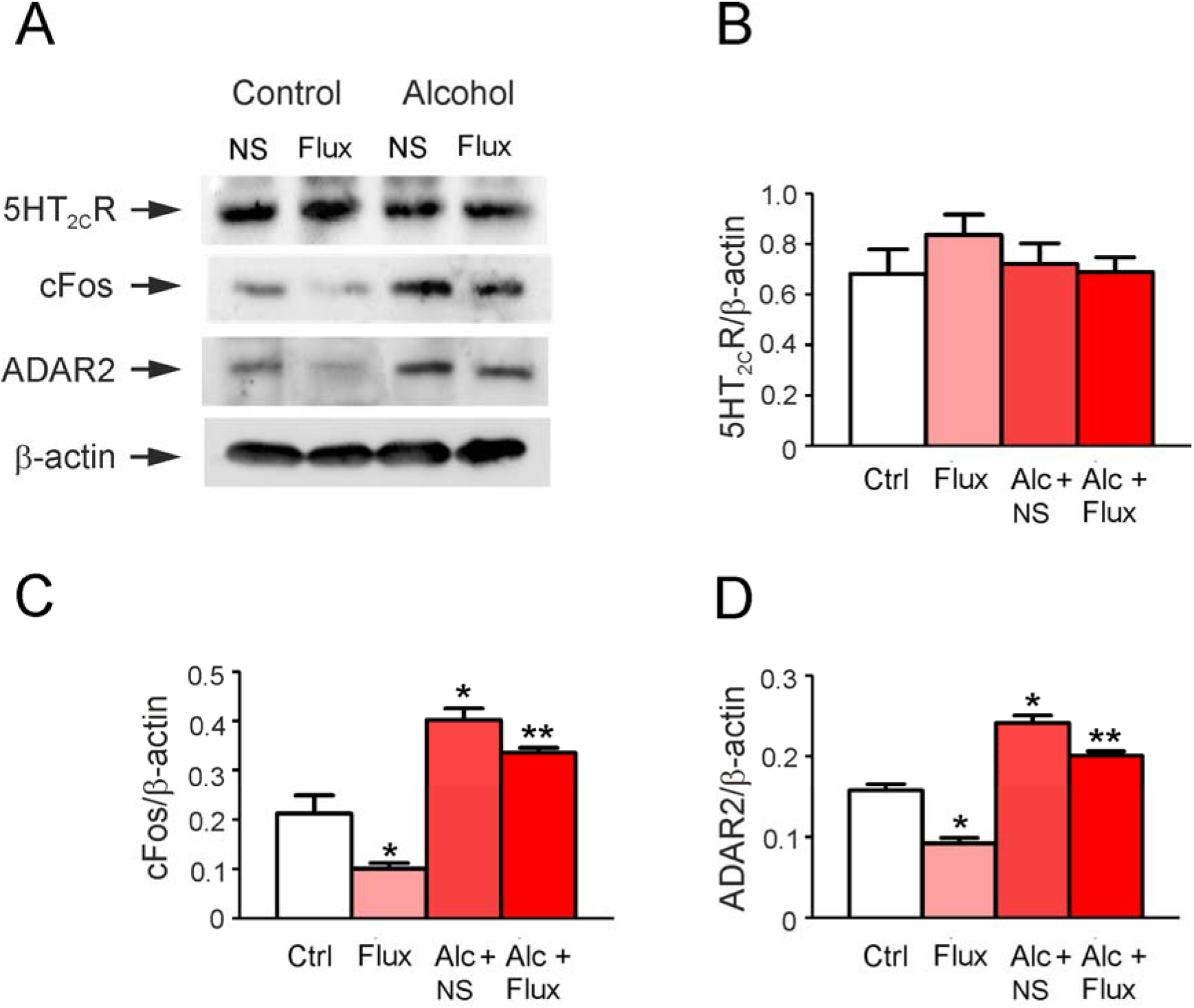
Fluoxetine suppressed the expression of cFos and ADAR2 increased by alcohol. Pre-treatedt with normal saline (NS; control) or with 4 mg/ml alcohol for 30 minutes, the primary cultures of astrocytes were exposed to NS or 1 μM fluoxetine for 24 hours. **(A)** Representative western blots showing protein levels of 5-HT_2C_R, cFos, ADAR2 and β-actin. Similar results were obtained from six independent experiments. Average expressive of 5-HT_2C_R, cFos and ADAR2 was quantified as ratio between 5-HT_2C_R and β-actin (**B**), ratio between cFos and β-actin (**C**), or ratio between ADAR2 and β-actin (**D**). Data represent mean ± SEM, n = 6. *Indicates statistically significant (p < 0.05) difference from any other group; **indicates statistically significant (p < 0.05) difference from control (Ctrl) and alcohol (Alc) groups.

Effects of fluoxetine on cFos and ADAR2 expression was antagonized by several inhibitors of cellular signalling cascades. As shown in Fig. 3B and 3C, SB242084 (SB), an inhibitor of 5-HT_2C_R, increased the expression of cFos and ADAR2 to 117.08 ± 10.18% (n = 6, p = 0.242) and 91.73 ± 7.97% (n = 6, p = 0.370) of control group. The PP1, a selective antagonist of Src, increased the level of cFos and ADAR2 to 114.78 ± 10.19% (n = 6, p = 0.271) and 88.08 ± 8.54% (n = 6, p = 0.320) of control group. The AG1478 (AG), a specific inhibitor of EGFR, enhanced the expression of cFos and ADAR2 to 130.04 ± 19.74% (n = 6, p = 0.189) and to 71.70 ± 3.28% (n = 6, p=0.130) of control group, but there was no significant difference between fluoxetine in combination with AG group and control (Ctrl) group regarding the expression of cFos or ADAR2. The LY294002 (LY), an inhibitor of AKT phosphorylation, changed the level of cFos and ADAR2 to 87.57 ± 15.93% (n = 6, p = 0.333) and 101.84 ± 5.78% (n = 6, p = 0.469) of control group.

**Figure 3.**
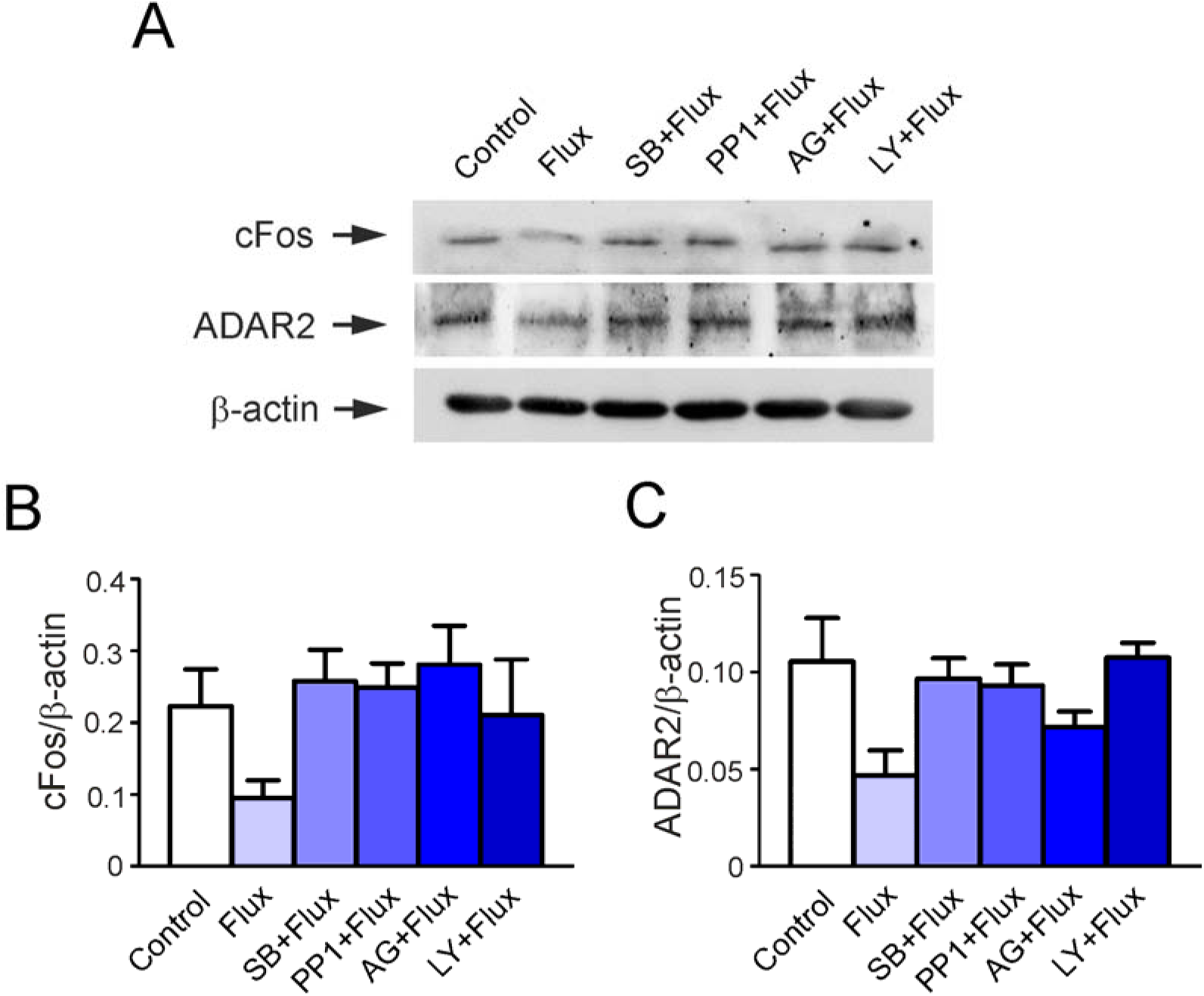
The regulation of fluoxetine in the expression of cFos and ADAR2. Pre-treatmed with normal saline, 400 nM SB204741 (SB; an inhibitor of 5-HT2BR), 10 μM PP1 (an inhibitor of Src), 10 μM AG1478 (AD; an inhibitor of EGFR), 10 μM LY294002 (LY; an inhibitor of p-AKT) for 30 minutes, primary cultured astrocytes were treated with NS or 1 μM fluoxetine for 24 hours. **(A)** Representative western blots protein levels of cFos, ADAR2 and β-actin. Similar results were obtained from six independent experiments. Average expression of cFos and ADAR2 was quantified as ratio between cFos and β-actin (**B**), or ratio between ADAR2 and β-actin (**C**). Data are shown as mean ± SEM, n=6. *Indicates statistically significant (p<0.05) difference from any other group.

### Alcohol and fluoxetine have opposite effects on RNA editing of 5-HT_2C_R

As shown in Fig. 4A and 4B, three subunits of ADAR catalyze editing of adenosine (A) to inosine (I) which is recognized as guanosine (G) at specific five sites of pre-mRNA (A-E). The ratios between the edited G-containing and unedited A-containing isoforms at site D of 5-HT_2C_R were determined. In the control group, RNA editing ratio between edited (G) and the sum of edited and unedited 5-HT_2C_R (G+A) was 77.08 ± 1.09% (n = 6). Fluoxetine decreased the ratio of G/G+A to 88.58 ± 1.48% of control group (n = 6, p = 0.001) at 1 μM, the treatment with alcohol increased the percentage of G/G+A to 104.54 ± 0.58% of control group (n = 6, p = 0.024) at 4 mg/ml. However, the administration of fluoxetine suppressed this editing ratio to 95.38 ± 0.79% of alcohol group (n = 6, p = 0.003), there was no significant difference between control group and alcohol plus fluoxetine group (Fig. 4D).

**Figure 4.**
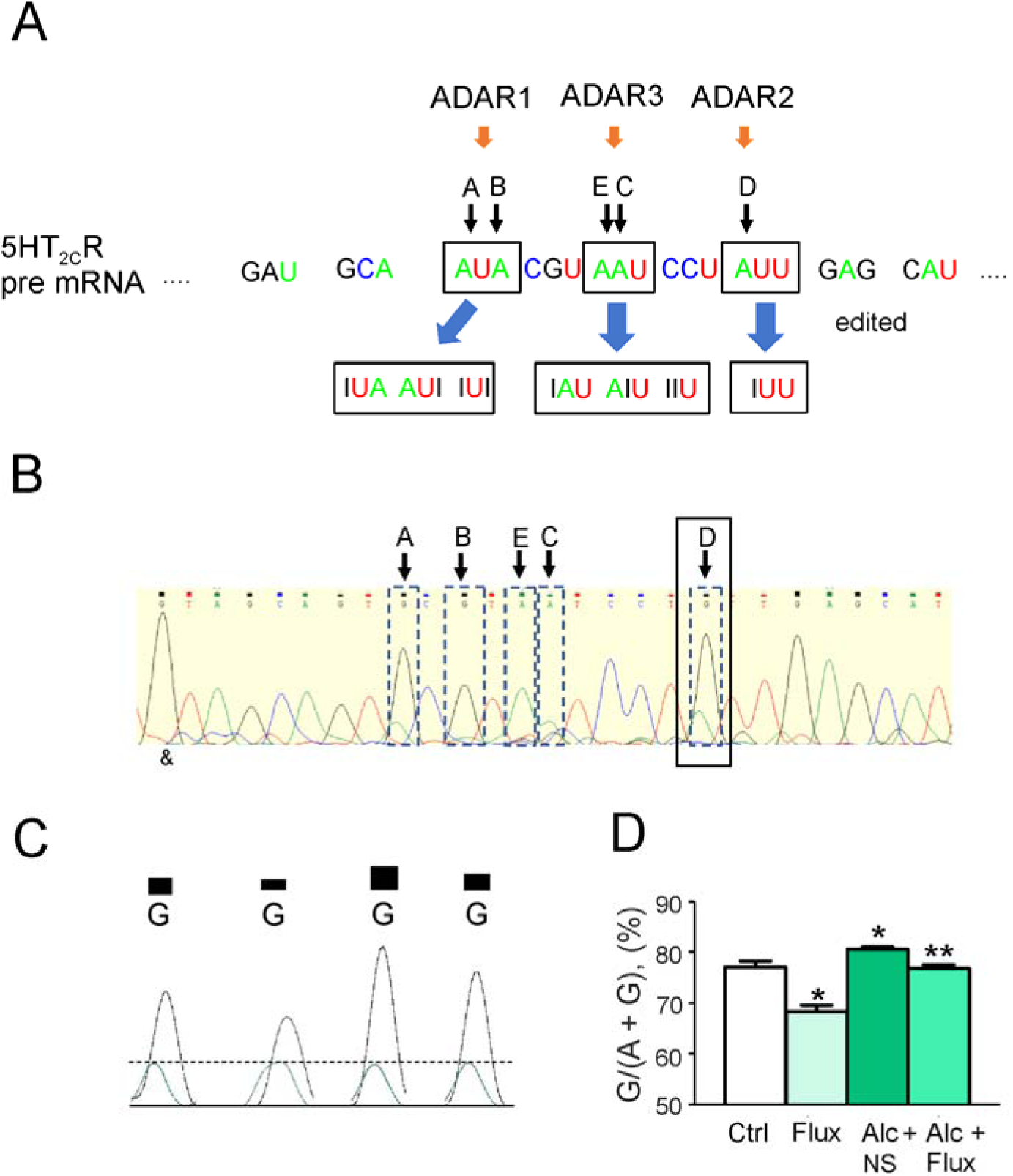
Fluoxetine decreased RNA editing percentage of 5-HT_2C_R enhanced by alcohol. The mice were feed with water or different concentration of alcohol for 36 days, then the mice were intraperitoneal injected (i.p.) with normal saline (NS) or fluoxetine (10 mg/kg/day) in the last week. (**A**) The pre-mRNA of 5-HT_2C_R can be edited by ADARs at five sites named A-E. (**B**) One representative mRNA sequencing data in control group. (C) The representative mRNA sequencing images at D site, the peak of adenosine (A) was green, the peak of guanosine (G) was black. (D) The ratio of G and G+A was shown mean ± SEM, n = 6. *Indicates statistically (p < 0.05) significant difference from control groups; **indicates statistically (p < 0.05) significant difference from alcohol (Alc) group.

**Figure 5.**
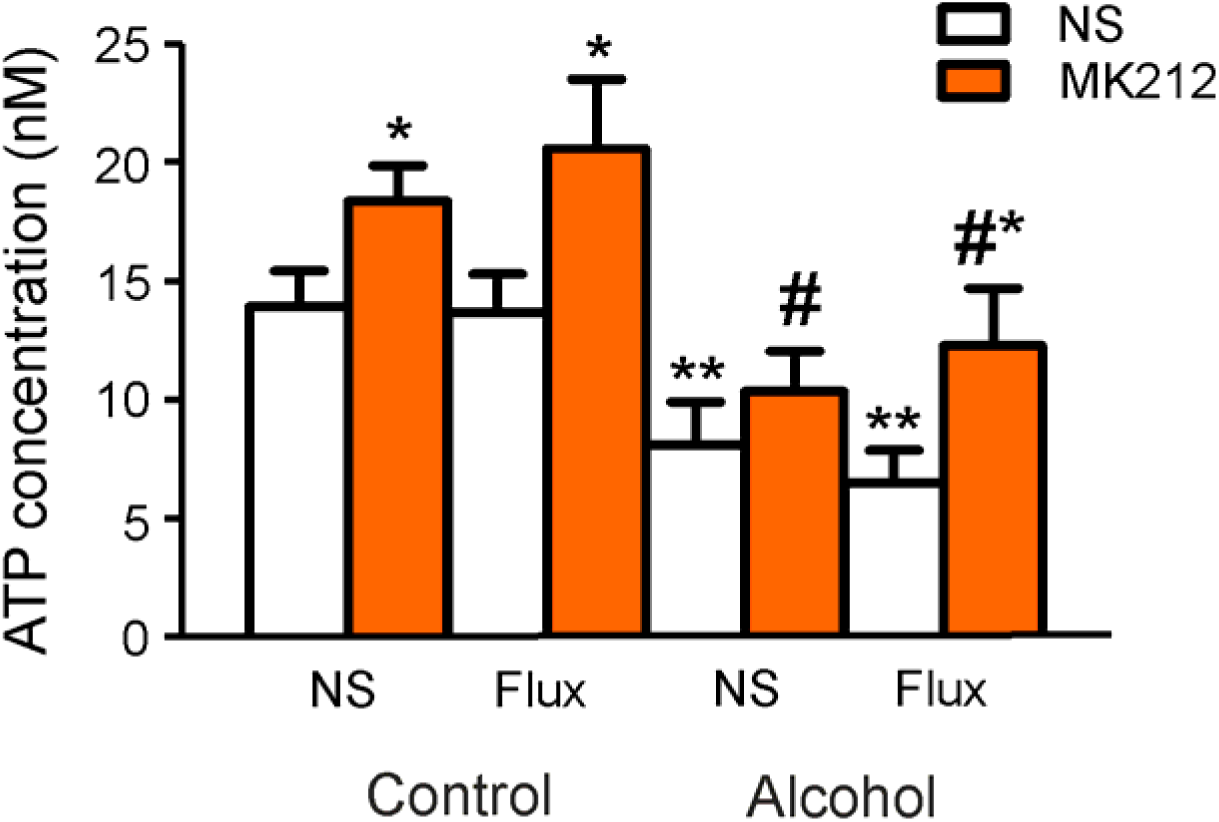
Fluoxetine enhanced the release of ATP decreased by alcohol via stimulating 5-HT_2C_R in astrocytes. Pre-treated with normal saline (NS; control) or 4 mg/ml alcohol for 30 minutes, cultured astrocytes were treated with NS or 1 μM fluoxetine for 24 h; extracellular ATP levels were measured after the administration of NS or MK212 (a selective agonist of 5-HT_2C_R) for 15 minutes. Data are shown as mean ± SEM, n = 6 *Indicates statistically significant (p<0.05) difference from NS group in the same group; **indicates statistically significant (p<0.05) difference from the four groups of control group; #indicates statistically significant (p<0.05) difference from two MK212 groups of control group.

### Alcohol and fluoxetine have opposite effect on 5-HT_2C_R-stimulated release of ATP

In the primary culture of astrocytes, MK212 (a selective agonist of 5-HT_2C_R) increased the level of extracellular ATP to 18.35 nM (n = 6, p = 0.046), which was 131.74 ± 13.93% of control group bathed normal saline (NS). After pre-incubation with 1 μM fluoxetine for 1 day, MK212 enhanced the level of ATP to 142.72 ± 16.31% of control group (n = 6, p = 0.039), but exposure of astrocytes to fluoxetine alone did not change extracellular ATP concentration, and there was also no significant difference between two MK212 group of NS and fluoxetine groups. In the presence of alcohol for 1 day, the extracellular level of ATP was decreased to 58.00 ± 11.40% (n = 6, p = 0.017), in the cells exposed to alcohol and fluoxetine ATP level was decreased to 46.64 ± 8.57% (n = 6, p = 0.003), compared with NS group without alcohol. Comparing with NS group treated with alcohol, the level of ATP induced by MK212 was increased to 128.85 ± 15.97% (n = 6, p = 0.192), there was no significant difference. After treatment with alcohol plus fluoxetine, MK212 however stimulated ATP release to 188.99 ± 29.25% of NS group (n = 6, p = 0.043).

### Fluoxetine improves alcohol-impaired behaviour

The gradually increased dose of alcohol was used for building an addiction model, whereas mice in control group were fed with water for 29 days (Fig. 6A). Subsequently “alcoholic” mice were divided into two groups, which were injected with NS or fluoxetine (10 mg/kg dose, i.p.) for one week; meanwhile treatment with alcohol was continued (Fig. 6A). Before the injection of fluoxetine, the ratio of alcohol preference was increased to 128.85 ± 5.98% of control group (n = 12, p = 0.011) on 29^th^ day (Fig. 6C). After the combined administration of alcohol with fluoxetine for one week, the preference ratio in alcohol plus fluoxetine group was decreased to 86.12 ± 2.17% of alcohol plus NS group (n = 6, p = 0.004), the alcohol plus NS group was increased to 156.74 ± 6.38% of control group (n = 6, p < 0.001) (Fig. 6D). In the open field test, compared with control group, the distance was increased to 130.42 ± 8.57% in alcohol plus NS group (n = 6, p = 0.019). However, this distance in alcohol plus fluoxetine was decreased to 82.82 ± 5.76% of alcohol plus NS group (n = 6, p = 0.045). As to another measured indicator in open field test, time spent in the centre was suppressed to 43.81 ± 7.14% of control group (n = 6, p = 0.007), fluoxetine increased this time to 98.07 ± 12.81% of control group (n = 6, p = 0.469). In the open field test, treatment with fluoxetine did not significantly change the travelled distance and staying time in centre area (Fig. 6F). For checking movement capacity by rotating rod test, alcohol reduced the time of mice standing on the rod to 30.32 ± 3.05% of control group (n = 6, p<0.001), the treatment with fluoxetine increased the dwell time reduced by alcohol to 60.65 ± 10.55% of control group (n = 6, p = 0.017) (Fig. 6G). Measuring mice depressive-like behaviour by tail suspension test, the recorded immobility time was increased by alcohol to 206.21 ± 16.11% of control group (n = 6, p = 0.002), fluoxetine decreased the immobility time to 68.45 ± 9.13% of alcohol group (n = 6, p = 0.012) (Fig. 6H). At the end of all tests, the weight of mice was also measured, there was no significant difference between groups (Fig. 6B).

**Figure 6.**
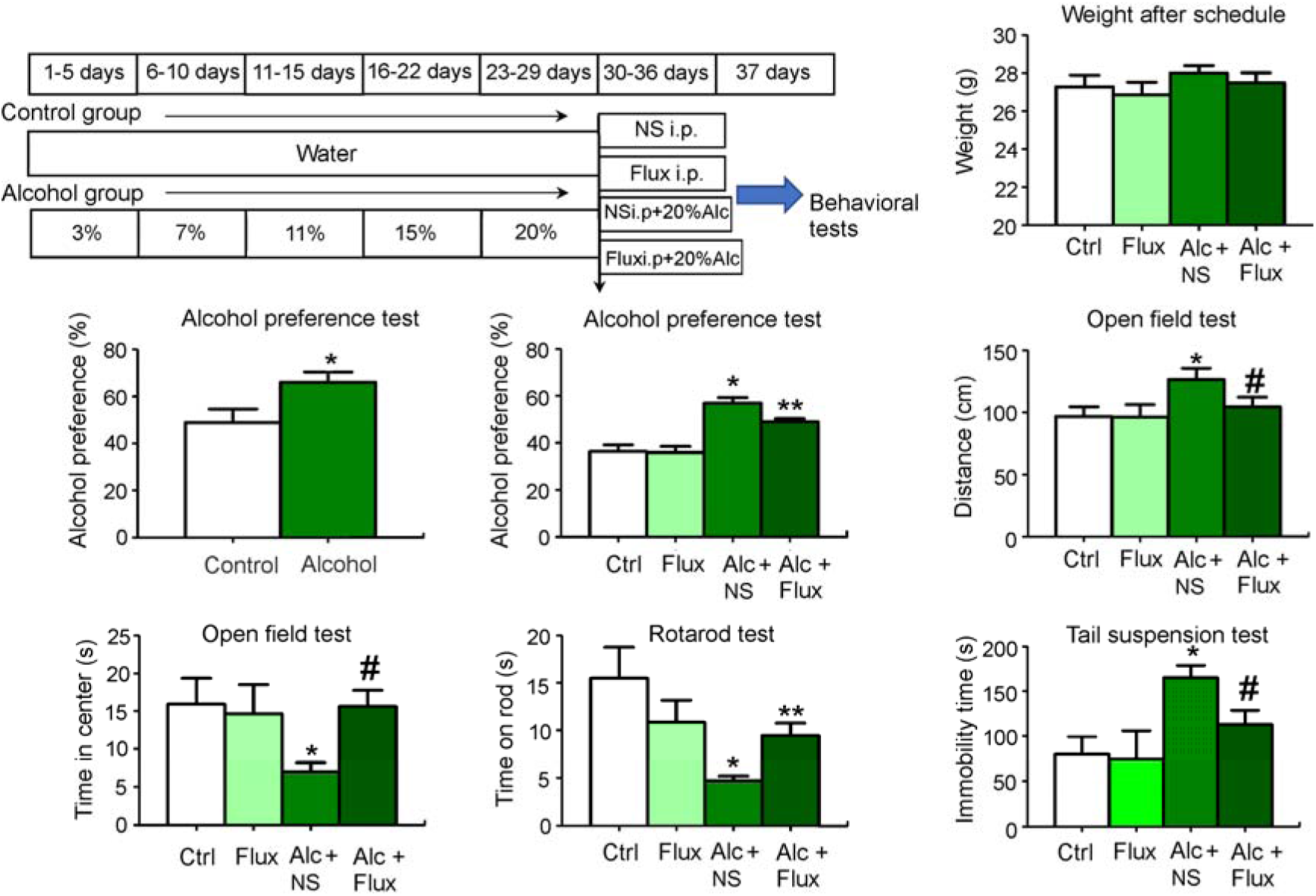
Fluoxetine improved the behaviour patterns induced by chronic alcohol addiction. **(A)** Mice were feed with water or different concentration of alcohol for 36 days, then the mice were intraperitoneal injected (i.p.) with normal saline (NS) or fluoxetine (10 mg/kg/day) in the last week. On the 30^th^ and 37^th^ day, alcohol preference test and behavioural measurements were performed. **(B)** The weight of mice was measured at the end of all treatments. Alcohol preference test was checked before the injection of fluoxetine (**C**) and after injection for one week (**D**). In the open field test, the travelled distance (**E**) and the time spent in the central area (**F**) were calculated. In rotarod test, the time dwelling on the rotarod was recorded **(G).** In tail suspension test, the time of immobility was determined **(H)**. Data are shown as mean ± SEM, n = 6. *Indicates statistically significant (p < 0.05) difference from control (Ctrl) group; **indicates statistically significant (p < 0.05) difference from Ctrl and alcohol (Alc) group; #indicates statistically significant (p < 0.05) difference from Alc group.

## Discussion

In this study, we discovered that the treatment with alcohol increased the expression of ADAR2, which promoted RNA editing of 5-HT_2C_R in astrocytes; an increased pool of edited 5-HT_2C_R inhibited the release of ATP induced by MK212. Fluoxetine at 1μM selectively stimulated 5-HT2BR and transactivated EGFR, which by means of Src, initiated PI3K/AKT pathway and down-regulated the expression of cFos and ADAR2, which ultimately suppressed RNA editing ratio of 5-HT_2C_R induced by alcohol and rescued the release of ATP stimulated by MK212. In experiments *in vivo*, in mouse alcohol addiction model, administration of fluoxetine decreased the alcohol preference, while simultaneously improving motor coordination, anxiety and despair-like behaviour (Fig. 7).

**Figure 7.**
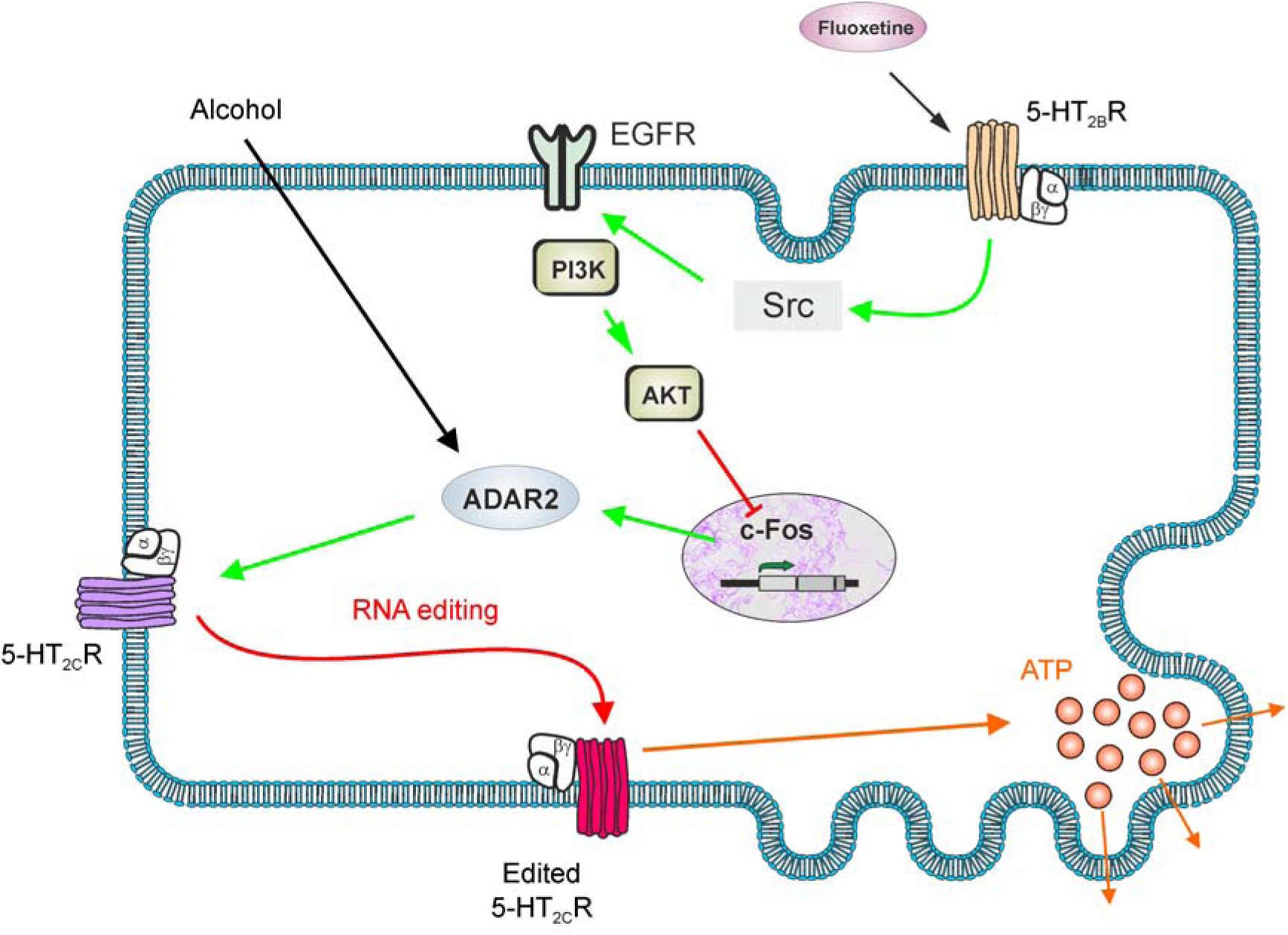
Fluoxetine alleviates alcohol effects by decreasing the expression of ADAR2 in astrocytes. The chronic alcohol addiction increased the expression of ADAR2, which stimulated editing of mRNA of 5-HT_2C_R and made this receptor dysfunctional. Stimulating 5-HT_2C_R with a selective agonist (MK212) increased the level of ATP released from astrocytes. Chronic alcohol treatment reduced the MK212-induced ATP release via upregulating the RNA editing effect of ADAR2. The decreased release of ATP from astrocytes could induce depressive like behaviours. However, fluoxetine at 1 μM transactivated Src-dependent EGFR and the downstream PI3K/AKT signalling pathway via stimulating 5-HT2BR, thereby inhibiting the expression of cFos and ADAR2. Fluoxetine decreased the RNA editing percentage of 5-HT_2C_R mRNA by suppressing the expression of ADAR2, hence recovering the function of 5-HT_2C_R impaired by alcohol. As a result, fluoxetine increased the MK212-induced release of ATP, which is decreased in chronic alcohol addiction. The recovery of ATP release from astrocytes may explain positive effect of fluoxetine on mice behavioural deficits triggered by chronic alcohol abuse.

Reduced release of ATP from astrocytes is related to depressive-like behaviors^20^. We found that alcohol decreased the release of ATP by up-regulating expression of ADAR2 which promoted RNA editing of 5-HT_2C_R thus rendered this receptor dysfunctional. Fluoxetine down-regulated the expression of ADAR2 and hence rescued ATP secretion by recovering the function of 5-HT_2C_R. The reduced ATP release from astrocytes may be due to the dysfunction of mitochondria. Alcohol withdrawal decreases the conversion of ADP to ATP in TCA cycle and inhibits ATP synthase in order to impede mitochondrial function via stimulating excitatory neurotransmitter glutamate in Purkinje cells^29, 30^. Moreover, the neurone-derived glutamate induced by stress triggers the release of ATP from ^31, 32^. However, the effect of chronic alcohol addiction on the regulation of ATP in neural cells is still unclear; although the decreased ATP level may contribute to the development of alcohol addiction.

RNA editing of 5-HT_2C_R via ADARs has been linked to alcohol preference^33^. Mice with specific ADAR2 deletion in nucleus accumbens were resistant to depressive or anxiety-like behaviours associating with low alcohol intake^34^. Administration of fluoxetine could decrease mRNA editing ratio of 5-HT_2C_R enhanced by alcohol by suppressing the expression of ADAR2 in astrocytes, thereby improving the behavioural phenotypes associated with chronic alcohol addiction in mice. In the present study we found that stimulation of 5-HT_2C_R using selective agonist increases the release of ATP from astrocytes. Arguably, stimulation of 5-HT_2C_R triggers the release of intracellular Ca^2+^ via PLC/IP3 signalling pathway thus stimulating secretion of ATP^35^. Alternatively activation of 5-HT_2C_Rs may recover mitochondrial function thus increasing the expression of ATP synthase and elevating the level of ATP, as was shown in NRK-52E cells^36^.

Treatment with various concentration of fluoxetine regulates the expression of cFos through the opposite effects of PI3K/AKT and MAPK/ERK signalling pathways in astrocytes. The cFos is a transcription factor which regulates expression of several genes, such as caveolin-1 and BDNF^15, 16, 37^. Here we found that fluoxetine at 1 μM suppressed the expression of ADAR2 increased by alcohol by regulating cFos through Src-mediated transactivation of EGFR and the downstream PI3K/AKT signalling pathway in astrocytes. Moreover, Src phosphorylation of EGFR at Y845 site is regulated by PI3K/AKT pathway^38^.

In conclusion, our results suggest that fluoxetine at low concentration has protective effects on the locomotor impairment and mental malfunction induced by alcoholism. Fluoxetine stimulates 5-HT_2C_R and the release of ATP via decreasing the RNA editing controlled by ADAR2 on astrocytes. In addition, fluoxetine is beneficial for suppressing the alcohol intake and preventing alcohol addiction.

## Acknowledgments

This study was supported by Grant No. 81871852 to BL from the National Natural Science Foundation of China, Grant No. XLYC1807137 to BL from LiaoNing Revitalization Talents Program, and Grant No. 20151098 to BL from the Scientific Research Foundation for Overseas Scholars of the Education Ministry of China. grant No. 81200935 to MX from the National Natural Science Foundation of China, Grant No. 20170541030 to MX from the Natural Science Foundation of Liaoning Province. Grant No. 81671862 and No. 81871529 to DG from the National Natural Science Foundation of China.

